# Using Natural Language Processing to Learn the Grammar of Glycans

**DOI:** 10.1101/2020.01.10.902114

**Authors:** Daniel Bojar, Diogo M. Camacho, James J. Collins

**Affiliations:** Wyss Institute for Biologically Inspired Engineering, Harvard University, Boston, MA 02115, USA; Department of Biological Engineering and Institute for Medical Engineering & Science, Massachusetts Institute of Technology, Cambridge, MA 02139, USA; Broad Institute of MIT and Harvard, Cambridge, MA 02142, USA

## Abstract

While nucleic acids and proteins receive ample attention, progress on understanding the structural and functional roles of carbohydrates has lagged behind. Here, we develop a language model for glycans, SweetTalk, taking into account glycan connectivity and composition. We use this model to investigate motifs in glycan substructures, classify them according to their O-/N-linkage, and predict their immunogenicity with an accuracy of ∼92%, opening up the potential for rational glycoengineering.

---

As one of life’s four fundamental polymers, glycans are involved in the pathophysiology of many critical diseases^1^. Glycans are the most abundant protein modification^2^, determining structure, stability, and function^3^; moreover, they have been identified on lipids^4^ and RNA^5^. Despite this, research on glycans has been slow. Recent advances in mass spectrometry-based glycomics profiling^6^ has sparked the systematic exploration of glycans and their overarching roles in biological processes. A myriad of issues, such as the lack of convenient bioinformatics resources for glycan sequences, the branched nature of glycans, and the extreme diversity of monosaccharides and bonds across species, has severely limited the computational study of glycans. Due to a near-complete lack of methodologies, existing analyses have predominantly focused on the protein associated with any given glycan rather than the carbohydrate chain itself^7^.

Machine learning and deep learning have recently entered the life sciences to help uncover highly nonlinear patterns, associations, and representations of biological data^8^. While many machine learning architectures could be envisioned for working with glycans, we chose a bidirectional recurrent neural network (RNN)^9^ as our model (Fig. 1a): (1) to capitalize on the sequential nature of glycans, (2) to learn a representation of glycans and their constituents, and (3) to construct a generative model for glycoengineering. We trained our model on a dataset of 21,296 glycans curated from a multitude of organisms, including humans and several model organisms (Supplementary Tables 1-2). To enable a contiguous language model despite the branched nature of glycans, we extracted partially overlapping “glycowords” from glycans (Supplementary Fig. 1), comprising three monosaccharides and two bonds, the largest substructure with strict linearity by definition. Using monosaccharides and bonds as “glycoletters”, we then trained a glycoletter-based language model, SweetTalk, with these glycowords (Supplementary Table 3), predicting the next most probable glycoletter given the preceding glycoletters. Model quality could be increased by pre-training a character-based language model (e.g., ‘G’, ‘a’, ‘l’) to construct initial glycoletter embeddings, which also led to more relevant glycoletter embeddings for downstream analyses (Supplementary Tables 4-5, Supplementary Fig. 2). An analysis of the learned embeddings of glycoletters after pre-training revealed similar positions in embedding space for monosaccharides and their modified counterparts (e.g., sulfurylated galactose, GalOS, and sulfurylated N-acetylgalactosamine, GalNAcOS, Fig. 1b, Supplementary Fig. 3). This result is reminiscent of observations made on the popular word2vec embeddings, in which semantically similar words form clusters^10^.

**Fig. 1:**
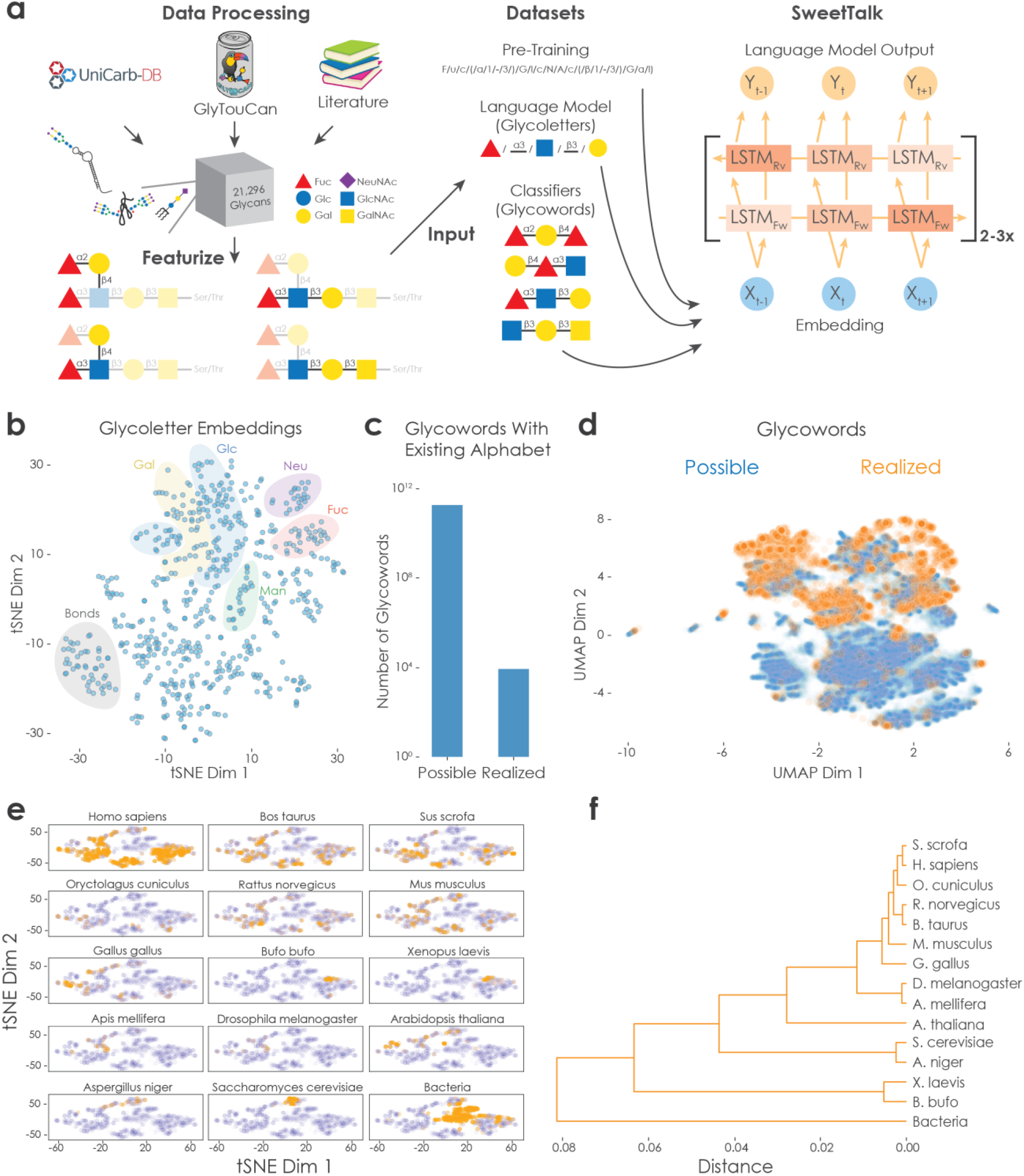
Learning the language of glycans revealed regularities in substructures and species distribution. **a**, Schematic of SweetTalk. Glycan sequences were extracted from UniCarbKB, GlyTouCan, and the academic literature, with the number of glycans after curation illustrated as a box. The diversity of glycans is represented by depicting exemplary glycans on proteins, lipids, and RNA. Branched glycan sequences were featurized by extracting glycowords, overlapping units consisting of three monosaccharides and two bonds. For the language models, these glycowords were used as input to train a character- or glycoletter-based bidirectional recurrent neural network predicting the next character or glycoletter. For the classifiers trained here, we used all glycowords of a given glycan as input. Both model types contained an initial embedding layer in which a representation of glycoletters and glycowords, respectively, could be learned from their context. Glycans are drawn in accordance with the symbol nomenclature for glycans (SNFG). **b**, Learned representation of glycoletters by SweetTalk. We used the trained embedding vector for every glycoletter to utilize its 128 dimensions as features for a t-distributed stochastic neighbor embedding (t-SNE). This allowed for the visualization of all observed glycoletters in embedding space. Areas enriched for modified monosaccharides of one type are colored. **c**, Comparing the abundance of possible and existing glycowords. Possible glycowords were calculated from the pool of observed glycoletters and their exhaustive combination (34 bonds and 554 monosaccharides). For the realized glycowords, all distinct glycowords in our dataset were counted. **d**, Comparing the distribution of possible and existing glycowords. Glycowords were generated by randomly sampling from the observed pool of monosaccharides and bonds. We repeated this process 250,000 times with replacement and discarded duplicate glycowords (though none were observed). Glycoword embeddings were formed by averaging their constituent glycoletter embeddings and these embeddings were used for a uniform manifold approximation and projection (UMAP) dimensionality reduction. Shown are 250,000 possible glycowords resulting from this process (blue) and all 8843 existing glycowords (orange). **e**, Species distribution of observed glycans in our dataset. For every glycan labeled with a species in our dataset (n = 2,481), we constructed an embedding by averaging their constituent glycoword embeddings. Subsequently, we used t-SNE to visualize these glycan embeddings in two dimensions. For every plot, glycans observed in a given species were colored in orange, while all other glycans were colored in blue. Note that several glycans were observed in multiple species. **f**, Hierarchical clustering dendrogram of species by their glycans. We averaged glycan embeddings from **e** for each species, constructed a cosine distance matrix between species, and performed hierarchical clustering on this distance matrix, visualized as a dendrogram.

**Fig. 2:**
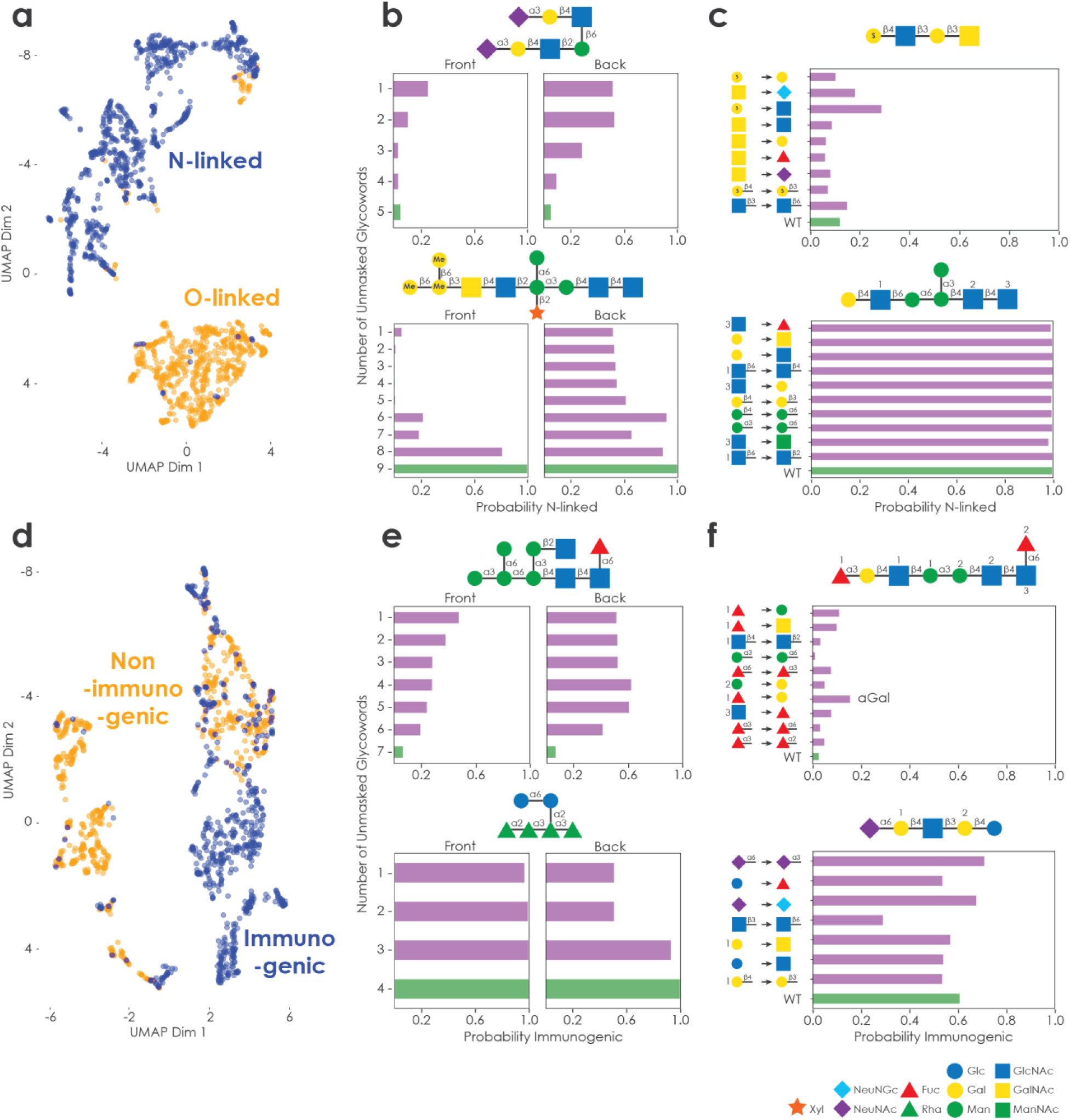
The glycan language model can be used to construct classifiers predicting O-/N-linkage and immunogenicity of glycans. **a**, Glycan embeddings learned by linkage classifier. Trained glycoword embeddings, extracted from our trained linkage classifier, were used to construct glycan embeddings via averaging of their constituent glycoword embeddings. After dimensionality reduction via UMAP, we plotted all glycans in our dataset with linkage information and colored them according to whether they were N-linked (blue) or O-linked (orange). **b**, Glycoword masking to probe linkage classifier. Glycowords were extracted from representative glycans and progressively exchanged with padding from both termini (‘masking’). ‘Front’ indicates the set of glycans in which all glycowords except the first were replaced by padding and then one glycoword after another was unmasked. Analogously, ‘Back’ depicts the progressive unmasking of glycowords starting from the last glycoword, and used as input for the trained linkage classifier. Inferred linkage probability is shown as a function of glycoword unmasking. The full-length glycan can be found at the bottom. **c**, Glycan *in silico* alterations to probe the linkage classifier. For 4000 iterations, single monosaccharides or bonds were replaced with a random monosaccharide or bond. The modified glycan was only retained and processed if the resulting glycowords existed in our dataset. Then, we used all modified glycans as input for the trained linkage classifier. Inferred linkage probability is plotted together with the altered glycan sequences, with the wildtype glycan found at the bottom. In case of ambiguity, a number indicates which monosaccharide was modified. **d**, Glycan embeddings learned by the immunogenicity classifier. Trained glycoword embeddings, extracted from our trained immunogenicity classifier, were used to construct glycan embeddings via averaging of their constituent glycoword embeddings. After dimensionality reduction via UMAP, we plotted all glycans in our dataset with immunogenicity information and colored them according to whether they were immunogenic (blue) or non-immunogenic (orange). **e**, Glycoword masking to probe the immunogenicity classifier. Glycowords were extracted from representative glycans, progressively exchanged with padding (‘masking’) from both termini (‘Front’/’Back’), and used as input for the trained immunogenicity classifier. Inferred immunogenicity probability is shown as a function of glycoword unmasking. The full-length glycan can be found at the bottom. **f**, Glycan *in silico* alterations to probe immunogenicity classifier. For 4000 iterations, single monosaccharides or bonds were replaced with a random monosaccharide or bond. The modified glycan was only retained and processed if the resulting glycowords existed in our dataset. Then, we used all modified glycans as input for the trained immunogenicity classifier. Inferred immunogenicity probability is plotted together with the altered glycan sequences, with the wildtype glycan found at the bottom. In case of ambiguity, a number indicates which monosaccharide was modified. All glycans are drawn in accordance with the symbol nomenclature for glycans (SNFG). The addition of an ‘S’ implies a sulfurylated monosaccharide, while ‘Me’ implies a methylated monosaccharide.

We then constructed initial glycoword embeddings by averaging their constituent glycoletter embeddings. Our first observation was that from the close to 200 billion possible glycowords (given the available glycoletters in our dataset) only 8,843 distinct glycowords (∼0.0000045%) were present in the glycans in our dataset (Fig. 1c). Moreover, these 8,843 glycowords were not evenly distributed in the learned embedding space, as existing glycowords formed clusters compared to generated, possible glycowords (Fig. 1d). The observation that the glycoword space (and, therefore, glycan space) is sparsely populated, led us to hypothesize that this is presumably a consequence of having to evolve dedicated enzymes for constructing specific glycan substructures. Building glycowords also allowed us to analyze their distribution in embedding space, uncovering, for instance, tight clustering of fucose- and rhamnose-containing glycowords, respectively (Supplementary Fig. 3). Constructing the corresponding embeddings for whole glycans allowed us to systematically investigate the distribution of glycans by species (Supplementary Table 6, Fig. 1e). Interestingly, glycans from *Saccharomyces cerevisiae*, known for their high mannose content^11^ (Supplementary Fig. 4), formed a tight cluster separate from most other species, except for the fungus *Aspergillus niger* which is also characterized by high-mannose glycans. Analogously, amphibian (*Xenopus laevis* and *Bufo bufo*) and insect species (*Drosophila melanogaster* and *Apis mellifera*), respectively, in our dataset also mostly formed their own clusters. Hierarchical clustering-based dendrograms using the glycan embeddings confirmed these observations despite different coverage of glycans between species (Fig. 1f), highlighting that there is preferential glycan usage within species.

Next, we used the glycoword embeddings as a starting point for an RNN-based classifier to predict the linkage of a given glycan to a protein, classifying the glycan as serine/threonine- (O-) or asparagine- (N-) linked (Fig. 2a, Supplementary Tables 7-8). Our model achieved a classification accuracy of >99% (F1 score: 0.994) on a dataset including fragmentary glycans, while the glycoword embeddings did not seem to change substantially during training (Supplementary Fig. 5), implying that the language model already extracted sufficient features for this task. Features, such as the near-perfect separation of galactose-, glucose-, and mannose-containing glycowords in the respective embedding space, were present for both the language model and the linkage classifier. By masking different parts of the glycans, we could then identify which regions of the glycan were most important for classification (Fig. 2b, Supplementary Fig. 6). These masking experiments demonstrated that the outer parts of glycans, in particular the outermost branching point residue, were more important for linkage classification. We performed *in silico* glycan alterations to probe further the classification determinants (Fig. 2c, Supplementary Fig. 7), discovering that particular glycans are more resilient to all allowed single glycoletter substitutions with regard to their predicted linkage. By restricting ourselves to alterations that resulted in existing glycowords, we further ensured that the resulting glycans are synthesizable by existing enzymes.

Given the important role glycans play in human immunity^12^, we curated a list of known immunogenic glycans from the literature, such as the ABH blood group antigens that are only immunoreactive in a certain fraction of the population as well as the yeast-derived O-mannosyl glycan or bacterial lipopolysaccharide constituents (Supplementary Table 9). We used our previously calculated glycoword embeddings to train an RNN-based classifier, allowing us to predict the immunogenicity of a glycan to humans purely from its raw sequence. On an independent validation dataset, to which the model had not been exposed before, our classifier achieved an accuracy of ∼92% (F1 score: 0.915), in what is the first-in-class algorithm for the classification of glycan immunogenicity (Fig. 2d, Supplementary Table 10). As the task of predicting glycan immunogenicity represents a substantially different task from the structural linkage prediction, we observed a shift in glycoword embeddings after training, distorting the previously observed clusters (Supplementary Fig. 8). Masking experiments again confirmed the importance of the more variable outer part of glycans for accurate classification (Fig. 2e, Supplementary Fig. 9). Notably, our trained classifier recognized the immunogenic potential of a recently discovered microbiome-produced inflammatory glycan (not present in our database) involved in Crohn’s disease^13^ (Fig. 2e, bottom). We also used our model to infer the human immunogenicity of porcine glycans, relevant for organ xenotransplantation and associated glycoengineering^16^, and identified glycans enriched for N-glycolylneuraminic acid (NeuNGc) or sulfurylated monosaccharides as potentially most immunogenic to humans (Supplementary Fig. 11, Supplementary Table 11).

Excitingly, *in silico* alteration experiments validated known immunogenic motifs, such as αGal^14^ (Gal(α1-3)Gal(β1-4)GlcNAc), which led to an increase in predicted immunogenicity (Fig. 2f, Supplementary Fig. 10). These developments could form the basis for a glycoengineering platform, as we have identified single modifications that abolished or induced immunogenicity, and could uncover novel immunogenic motifs. Additionally, we argue that modifications for increased immunogenicity could be critical for novel vaccine design, supported, in part, by the observation that broadly neutralizing antibodies in HIV-infected patients predominantly target glycans on the surface of HIV^15^. Bioengineering strategies using existing biosynthetic enzymes should be feasible by design, as we enforced all modified glycowords to correspond to existing glycowords.

Substantial efforts are currently being invested into glycoengineering for improved biopharmaceutical stability and effectiveness^17^. Additionally, the relevance of glycans for diseases ranging from inflammatory bowel disease^18^ to cancer^19^ has become increasingly apparent. The need for computational methods for glycan analysis has thus become even more pressing. Recent deep learning-based language models for DNA and RNA^20^, as well as proteins^21^, have enabled rational modifications and created new means to augment our understanding of these biological polymers. SweetTalk, our natural language model for glycans, contributes to the advancement of predictive glycoengineering and enables deeper exploration of glycobiology as a whole. The deep learning modeling strategies presented here allowed us to introduce a language model for glycans, while our curated dataset introduces a state-of-the-art coverage for glycan sequences across a multitude of organisms. In contrast to a word2vec-type model, our language model-based approach is able to capture sequential information beyond mere co-occurrences in glycan sequences. Additionally, starting from a glycoletter-based model allows for the construction of embeddings for close to 200 billion glycowords, making SweetTalk easily extendable. Our character-based language model even enables the construction of embeddings for novel glycoletters (Supplementary Fig. 12), further expanding the reach of our hierarchical approach.

The language model introduced here shows promise for the expansion of glycoengineering, molecular diagnostics, therapeutics, and general aspects of glycobiology, such as glycan evolution and preferential utilization across species. To the best of our knowledge, our curated and processed dataset of glycan sequences, including labels for linkages and species of origin, represents the largest dataset of its type, easily accessible to machine learning techniques. With this, we also include an expanded processed dataset containing 2,615 glycans that consisted of less than one glycoword and were too short for our analyses (Supplementary Table 12); we envision that these open-access resources will facilitate advances in glycobiology and beyond.

## Methods

All methods can be found in the online Methods section associated with this manuscript.

## Data availability

Data used for all analyses can be found in the supplementary tables.

## Code availability

All code and trained models can be found at https://github.com/midas-wyss/sweettalk.

## Acknowledgements

The authors would like to thank Xiao Tan, Jacqueline Valeri, Pradeep Ramesh, and Ashty Karim for helpful discussions.

## Contributions

D.B. conceived the method. D.B., D.M.C., and J.J.C. designed the experiments. D.B. performed the experiments and implemented the method. D.M.C. and J.J.C. supervised the work. D.B., D.M.C., and J.J.C. wrote and approved the manuscript.

