## Supplementary Information for "Using Natural Language Processing to Learn the Grammar of Glycans"

### Methods

#### Datasets

For the language model dataset, we retrieved all entries on UniCarbKB[1] that had a working link to PubChem[2] and contained at least three monosaccharides; we processed and extracted the glycan sequences, including their linkages (n = 1,687). We also extracted the Web3 Unique Representation of Carbohydrate Structures (WURCS) representation[3] of the set of all glycans deposited on GlyTouCan[4], which were also deposited on PubChem (n = 21,485). From the latter, we only processed glycans containing at least three monosaccharides for the language model dataset (n = 18,926). Glycans in WURCS representation were then reformatted into the IUPACcondensed representation, using the GlycanFormatConverter[5], for further processing. For the immunogenicity classifier, all GlycoEpitope (https://www.glycoepitope.jp) entries with a minimum length of at least three monosaccharides were extracted. This list was further complemented by targeted literature searches[6–16], resulting in the final set of immunogenic glycans (n = 685). This was combined into the final dataset used for training the language model (n = 21,296).

Analogous to word stemming in natural language processing, we curated this dataset to achieve an appropriate level of granularity, given our dataset size. We removed position-specific information of monosaccharide modifications, reporting only the type of monosaccharide modification. While this procedure created duplicate glycans in our dataset, we retained all glycans to preserve the relative proportion of glycoword-contexts for an appropriate weighting in our language model. Further modifications involved harmonization of capitalizing monosaccharide names and removal of dangling linkages (e.g., '(a1-'). For glycan repeat structures, the first monosaccharide was additionally appended to the end of the repeat structure to capture more glycoword contexts. To avoid isomorphic duplicates, we also enforced an order of monosaccharide modifications: NAc > OAc > NGc > OGc > NS > OS > NP > OP > NAm > OAm > NBut > OBut > NProp > OProp > NMe > OMe > CMe > NFo > OFo > OPPEtn > OPEtn > OEtn > A > N > SH > rest. The raw glycan sequences can be found in Supplementary Table 1, while the processed glycan sequences are provided in Supplementary Tables 2 and 11. Abbreviations for all monosaccharides and their modifications can be found in Supplementary Table 13.

Glycowords were then generated from all glycans, using a script extracting the longest possible strictly linear glycan substructures, consisting of three monosaccharides and two bonds. These glycowords were extracted with maximum overlap so that two subsequent glycowords only differed in one monosaccharide and one bond. The aim of these glycowords is to capture representative characteristics and local structural contexts of a given glycan. For language model pre-training, we split all monosaccharides and bonds of a glycoword into individual characters ('G', 'a', 'l', etc.) to pre-train our language model. The dataset comprising all glycowords (n = 113,112) was then used to train a context-specific, glycoletter-based language model.

The protein linkage classifier used entries from UniCarbKB, that included linkage information and corresponded to O- or N-linked glycans (n = 1,608, excluding free oligosaccharides). The immunogenicity classifier used immunogenic glycans from GlycoEpitope and the academic literature (n = 685), complemented with 685 randomly chosen human glycans from the language model dataset as negative examples. All sequences were processed identically to the language model dataset.

All entries corresponding to *Homo sapiens*, *Bos taurus*, *Sus scrofa*, *Oryctolagus cuniculus*, *Rattus norvegicus*, *Mus musculus*, *Gallus gallus*, *Bufo bufo*, *Xenopus laevis*, *Apis mellifera*, *Drosophila melanogaster*, *Caenorhabditis elegans*, *Arabidopsis thaliana*, *Aspergillus niger*, *Saccharomyces cerevisiae* on UniCarbKB and core oligosaccharide as well as O-antigen portions of lipopolysaccharide (LPS) of various bacteria, were used to analyze glycan distribution across species.

**Model training**

PyTorch[17] on a single NVIDIA® Tesla® K80 GPU was used to train all models reported here. For all models, architecture and hyperparameters were optimized by minimizing the respective loss function. We used mixed precision training utilizing the Apex library (https://github.com/nvidia/apex) for the language models. For the language model, the glycoword dataset was randomly split into training (80%, 90,490 glycowords) and validation sets (20%, 22,622 glycowords). For both classifiers, the dataset was split at the glycan level with the same proportions as for the language model. Additionally, glycans represented as a list of glycowords were brought to the same length by padding. Batch sizes were 256 and 64 for the training and validation sets of the language models, respectively. For the classifiers, we used a batch size of 32 for both training and validation.

SweetTalk consisted of a 128-dimensional embedding layer for all observed glycoletters (i.e., monosaccharides and bonds in glycans present in our datasets) with a subsequent two-layered (three-layered for classifiers), bidirectional LSTM (long short-term memory[18]) containing 128 nodes per layer. To learn a glycoletter-based language model, the next glycoletter was predicted for each glycoword. Therefore, the concatenated hidden representation learned by the bidirectional LSTMs was projected, via a fully connected layer, to a vector corresponding to the number of glycoletters. Fully connected layers were initialized by Xavier initialization[19]. Utilizing the ADAM optimizer[20] with a starting learning rate of 0.005 (language model pre-training), 0.001 (language model), or 0.0005 (both classifiers), we used a cosine function-based learning rate decay over the course of 50 to 150 epochs. The classification models also used a weight decay of 0.001 (linkage) or 0.005 (immunogenicity). An early stopping criterion was employed for regularization to cease training once the cross-entropy validation loss did not decrease for more than 10 epochs. For the classifiers, binary cross-entropy was used as the loss function.

**Hierarchical clustering dendrogram analysis**

Glycan embeddings, formed by averaging their constituent glycoword embeddings, were averaged for each species. Then, a cosine distance matrix was calculated using the SciPy library[21]. We used this distance matrix to perform hierarchical clustering via SciPy. The dendrogram was then created from this hierarchical clustering analysis.

**Generating random glycowords**

Random glycowords were created by sampling glycoletters for sequences corresponding to maximum-length linear glycowords (i.e., three monosaccharides connected by two bonds). At each position, either a monosaccharide or a bond was randomly chosen with replacement from the respective pool. Then, the generated glycoword was queried against the vocabulary of actually occurring glycowords to determine whether it corresponded to an observed glycoword. Duplicate glycowords were discarded.

### Masking and modification

For masking, glycowords in the respective glycans were incrementally replaced by padding. This was performed from both ends of glycans to identify position-specific effects on model performance. Masked glycans were used as inputs to the trained classifier, and the predicted class probability was compared to the full, unmasked glycan. For modifying glycans, glycoletters in the respective glycans were randomly exchanged with other glycoletters so that the resulting glycowords were still present in the vocabulary contained in our datasets. Then, altered glycans were utilized to compare class probabilities with the wildtype glycan using the trained classifier.

### Out-of-sample glycans

To enable predictions with our trained classifiers on glycans with unobserved glycowords, we needed to construct labels for the unobserved glycowords in order to pass the list of glycowords to the model for inference. All glycowords existing in our classifier embedding were converted to their respective labels. For glycowords not present in our embedding, we constructed an averaged embedding from our glycoletter-based language model. Then, we calculated the cosine similarity to all glycoword embeddings present in our trained classifier. Subsequently, we replaced the unobserved glycoword with the label corresponding to the observed glycoword with the highest cosine similarity.

### Supplementary Figures

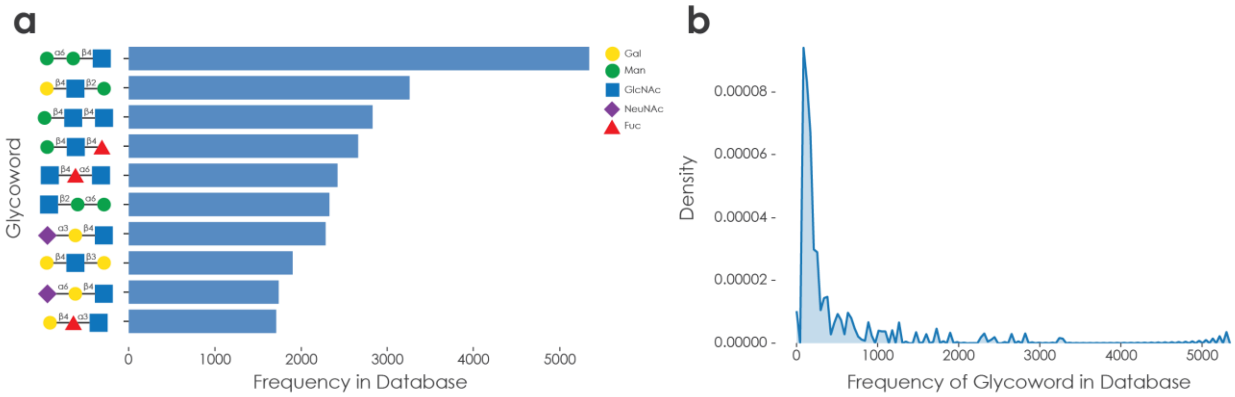


**Supplementary Fig. 1: Glycoword distribution in our database.** (**a**) Glycowords in our database were counted and the ten most frequent are shown with their abundance in all analyzed glycans. Glycans are drawn in accordance with the symbol nomenclature for glycans (SNFG). (**b**) For all observed glycowords, their frequency was determined and is depicted as a kernel density estimate plot, revealing a nearly bimodal distribution with many rare glycowords and a few extremely prevalent glycowords, depicted in (a).

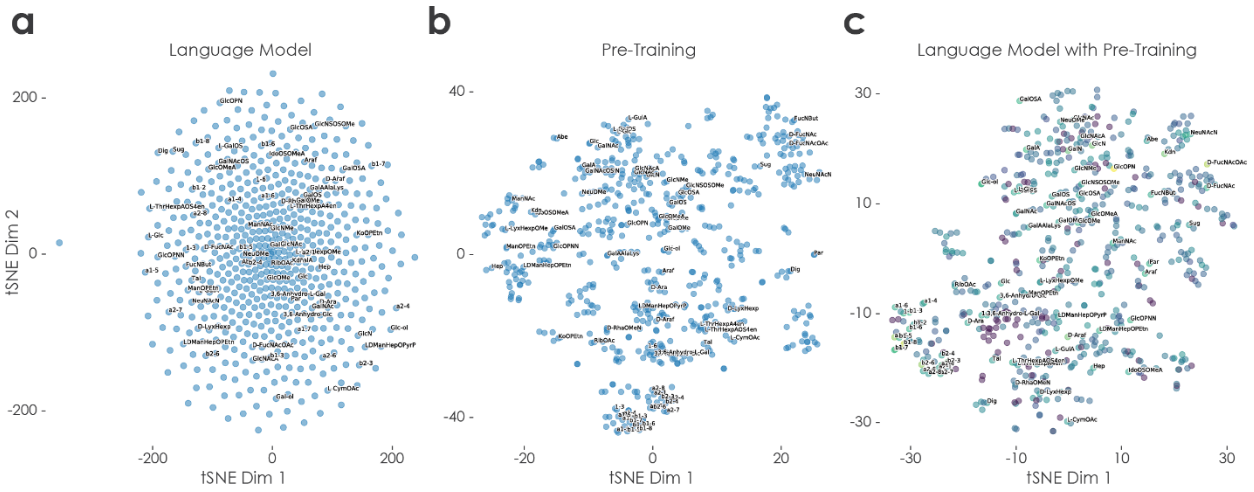


**Supplementary Fig. 2: Learned glycoletter representation for different language models.** We used the 128 dimensions of trained embedding vectors for every glycoletter as features for a t-distributed stochastic neighbor embedding (t-SNE). This allowed for the visualization of all observed glycoletters in embedding space. The depicted embeddings stem from: (**a**) a naively trained glycoletter-based language model, (**b**) a character-based pre-training, and (**c**) a glycoletter-based language model initialized with pre-trained glycoletter embeddings. Glycoletters in (c) are colored by Euclidean distance between their embeddings in (b) and (c). Glycoletters with at least 50% of the maximum Euclidean distance are shown with their names.

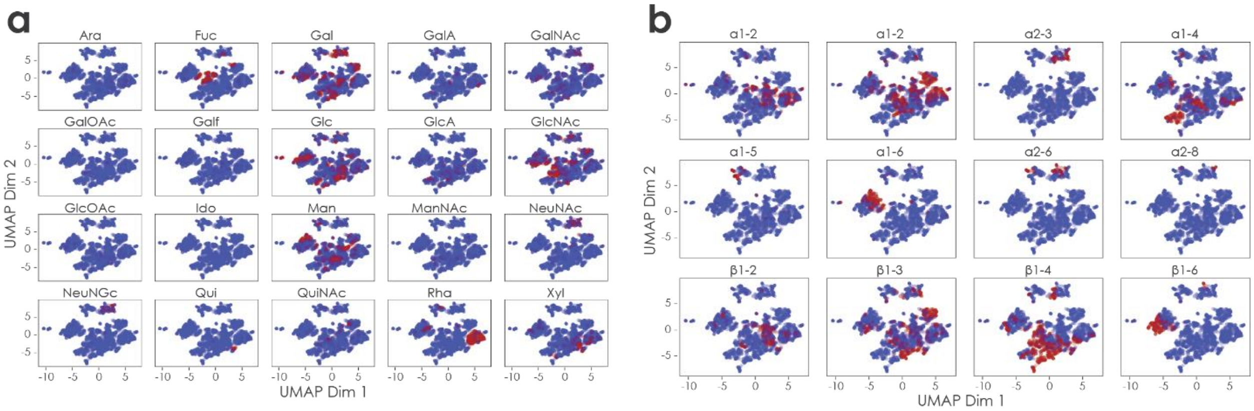


**Supplementary Fig. 3: Glycoword embeddings learned by language model.** Glycoword embeddings were constructed by averaging the learned embeddings of their constituent glycoletters after training SweetTalk. All 8,843 unique glycoword embeddings in our dataset were then dimensionality reduced via UMAP and are shown here. Further, glycowords were colored red if they contained one of a set of (**a**) common monosaccharides or (**b**) bonds.

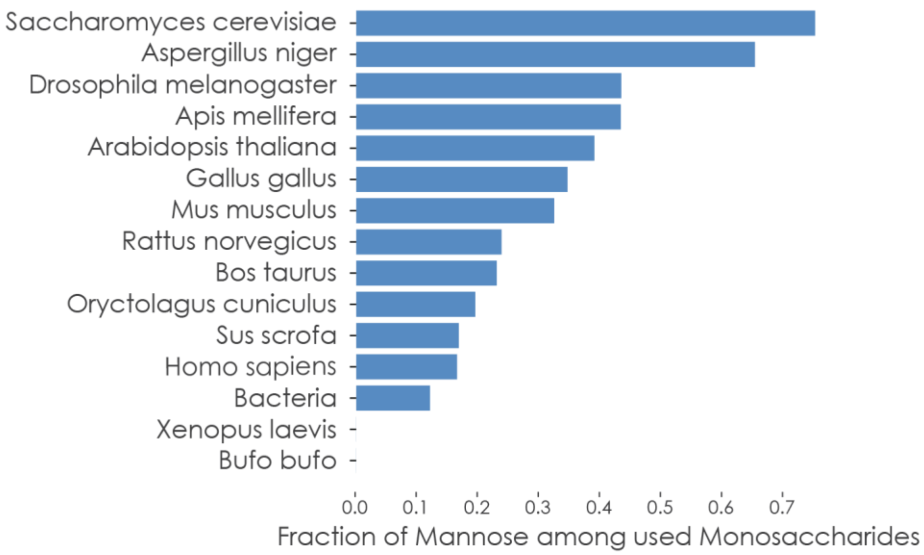


**Supplementary Fig. 4: Mannose content of glycans by species.** For each species, we counted the occurrence of mannose residues in each glycan and normalized this number by the total amount of monosaccharides in that glycan. The averaged fractions of mannose content in glycans (from 0 to 1) are plotted here for the described species.

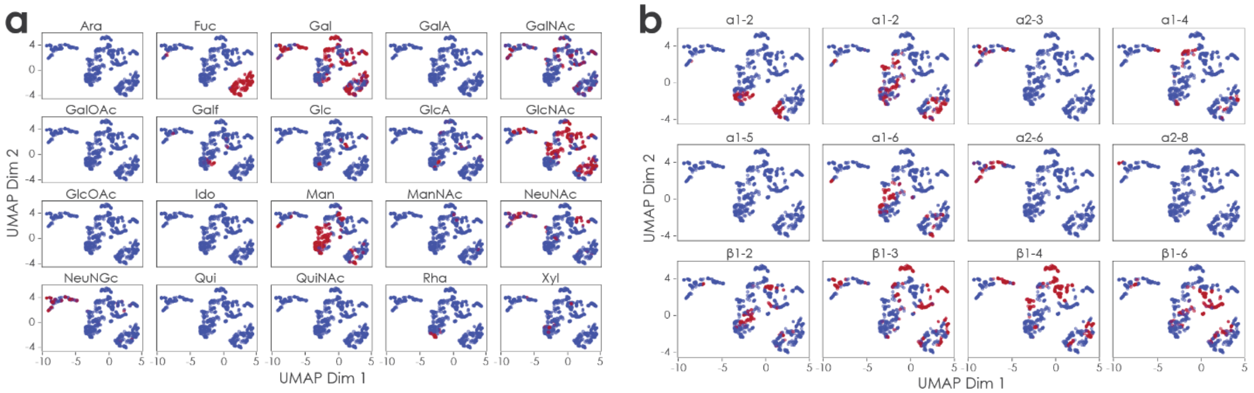


**Supplementary Fig. 5: Glycoword embeddings learned by linkage classifier.** Learned glycoword embeddings were extracted from the embedding layer of the linkage classifier. Embeddings of the 735 unique glycowords observed in our linkage dataset were then dimensionality reduced via UMAP and are shown here. Further, glycowords were colored red if they contained one of a set of common (**a**) monosaccharides or (**b**) bonds.

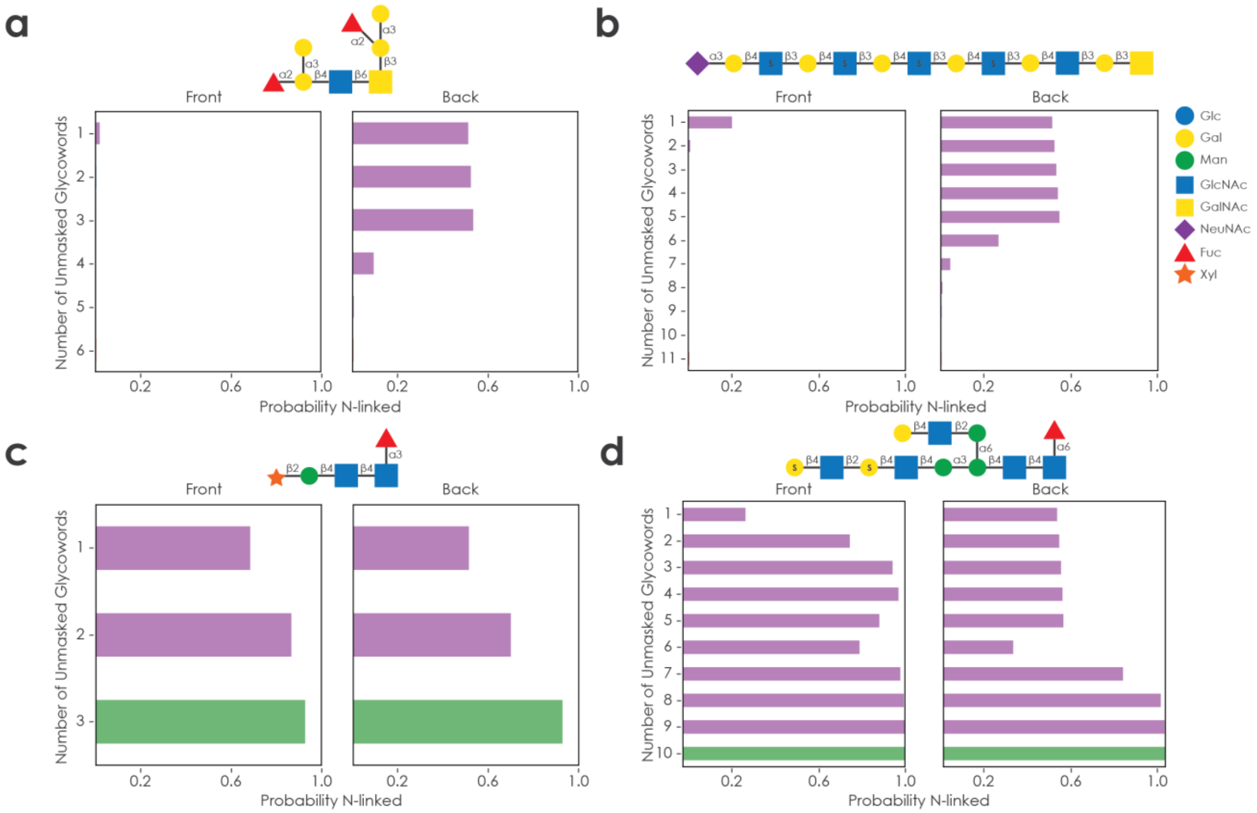


**Supplementary Fig. 6: Glycoword masking to probe linkage classifier.** (**a-d**) Representative glycans were processed into their glycowords. Glycowords were then progressively replaced with padding from both sides of the glycans. ‘Front’ indicates the set of glycans in which all glycowords except the first were replaced by padding and then one glycoword after another was unmasked. Analogously, ‘Back’ depicts the progressive unmasking of glycowords starting from the last glycoword. All glycan variants were then used with the trained linkage classifier for inference; the probability of the glycan being N-linked, as inferred by the model, is plotted here. By definition, a probability lower than 0.5 implies that the model predicted the glycan to be O-linked. The full-length glycan, with the correct prediction, is always located at the bottom. Glycans are drawn in accordance with the symbol nomenclature for glycans (SNFG). The addition of an ‘S’ implies a sulfurylated monosaccharide.

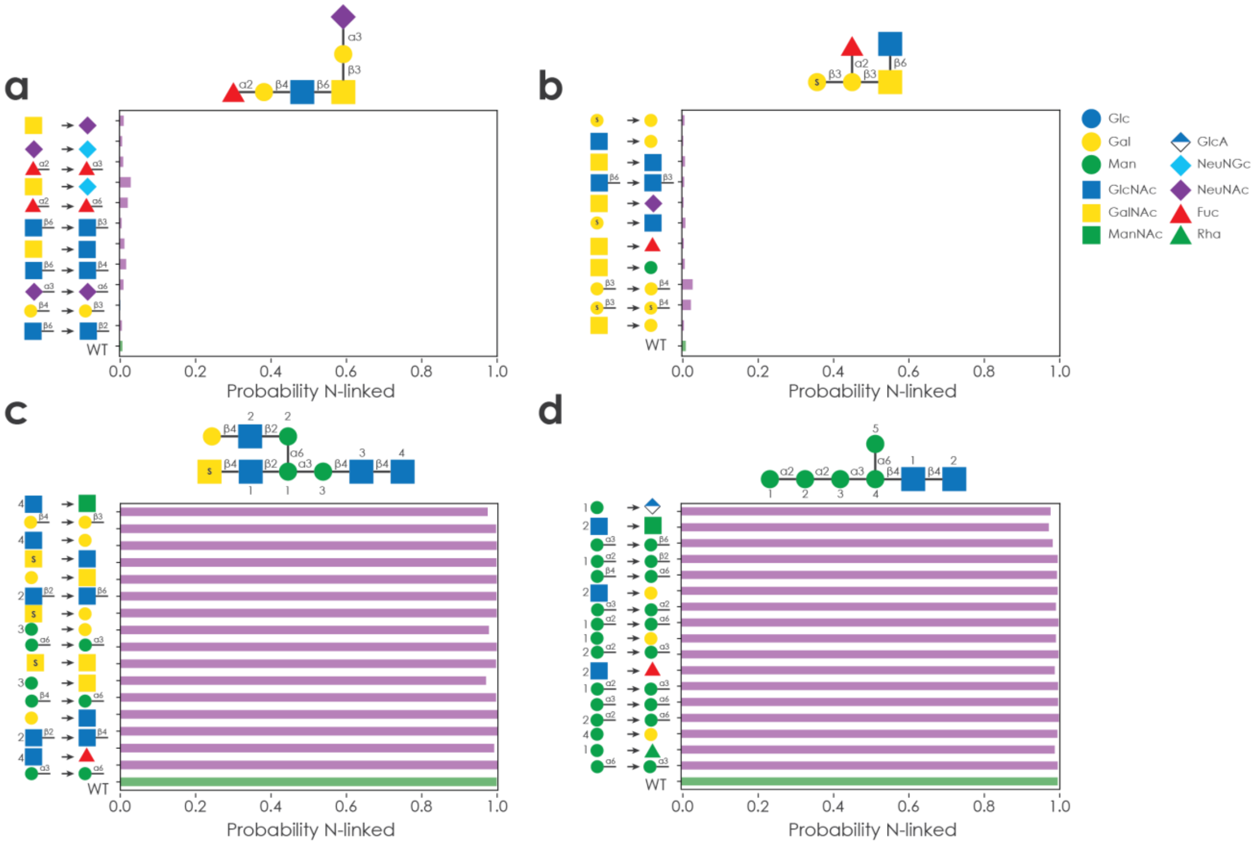


**Supplementary Fig. 7: Glycan *in silico* modification to probe linkage classifier.** (**a-d**) For 4000 iterations, a random position in a glycan was exchanged with a random monosaccharide or bond (depending on the position). The altered glycan was only retained and processed if the resulting glycowords existed in our dataset. Then, analogous to the masking experiments in Supplementary Fig. 6, we used our trained linkage classifier for inference and plotted the predicted probability of the modified glycans to be N-linked, with the wildtype glycan always located at the bottom. In case of ambiguity, a number indicates which monosaccharide was modified. Glycans are drawn in accordance with the symbol nomenclature for glycans (SNFG). The addition of an ‘S’ implies a sulfurylated monosaccharide.

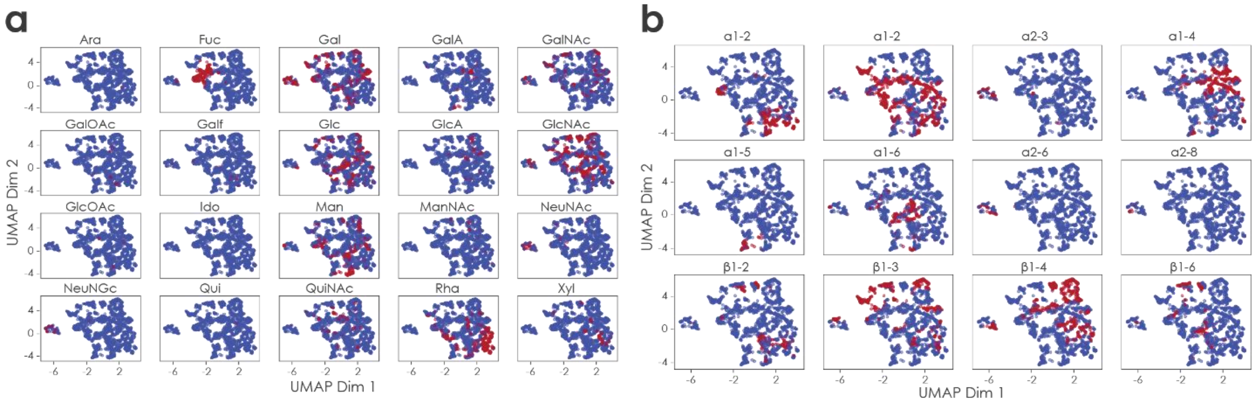


**Supplementary Fig. 8: Glycoword embeddings learned by immunogenicity classifier.** Learned glycoword embeddings were extracted from the embedding layer of the immunogenicity classifier. Embeddings of the 1,774 unique glycowords observed in our immunogenicity dataset were then dimensionality reduced via UMAP and are shown here. Further, glycowords were colored red if they contained one of a set of (**a**) common monosaccharides or (**b**) bonds.

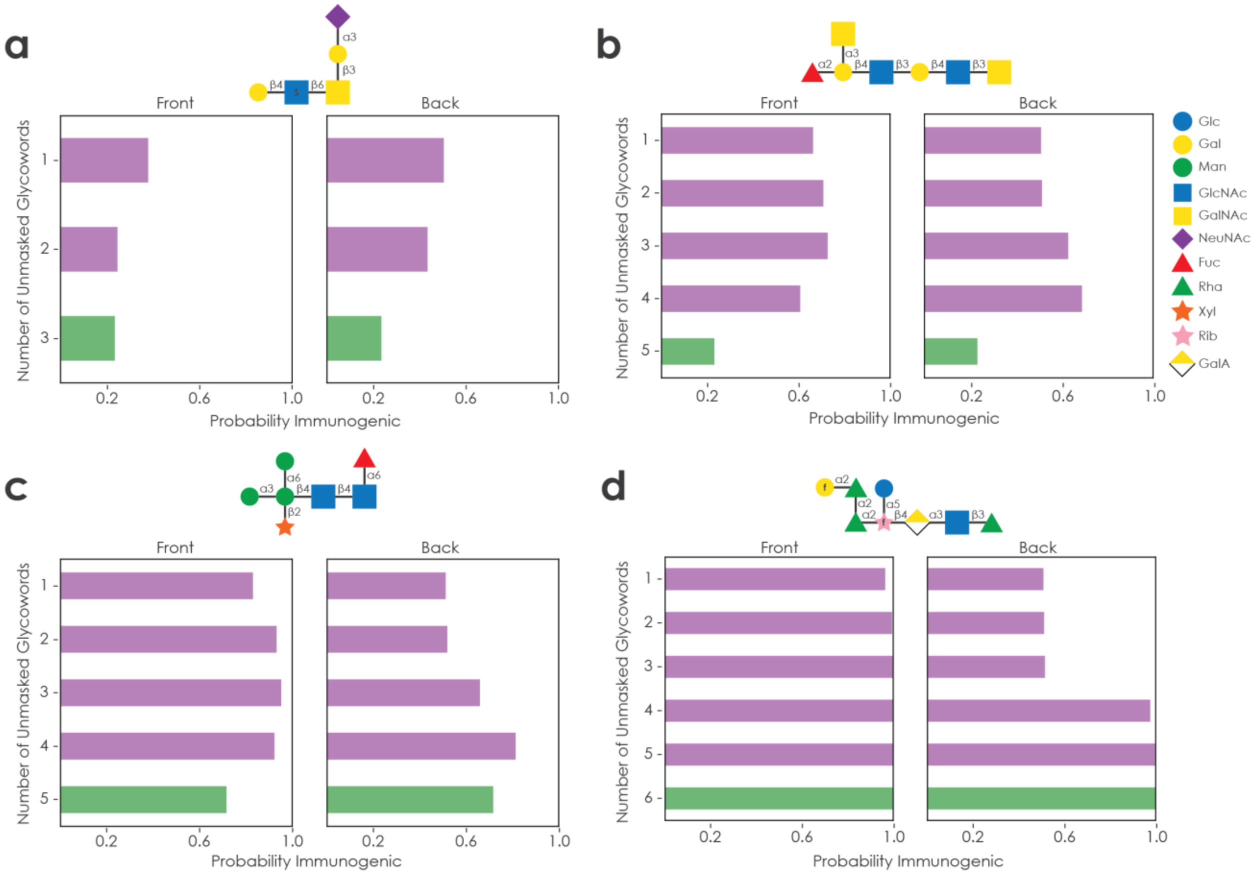


**Supplementary Fig. 9: Glycoword masking to probe immunogenicity classifier.** (**a-d**) Representative glycans were processed into their glycowords. Glycowords were then progressively replaced with padding from both sides of the glycans. 'Front' indicates the set of glycans in which all glycowords except the first were replaced by padding and then one glycoword after another was unmasked. Analogously, 'Back' depicts the progressive unmasking of glycowords starting from the last glycoword. All glycan variants were then used with the trained immunogenicity classifier for inference; the probability of the glycan being immunogenic, as inferred by the model, is plotted here. By convention, a probability lower than 0.5 implies that the model predicted the glycan to be non-immunogenic. The full-length glycan, with the correct prediction, is always located at the bottom. Glycans are drawn in accordance with the symbol nomenclature for glycans (SNFG). The addition of an 'S' implies a sulfurylated monosaccharide, while 'f' indicates the furanose form of the respective monosaccharide.

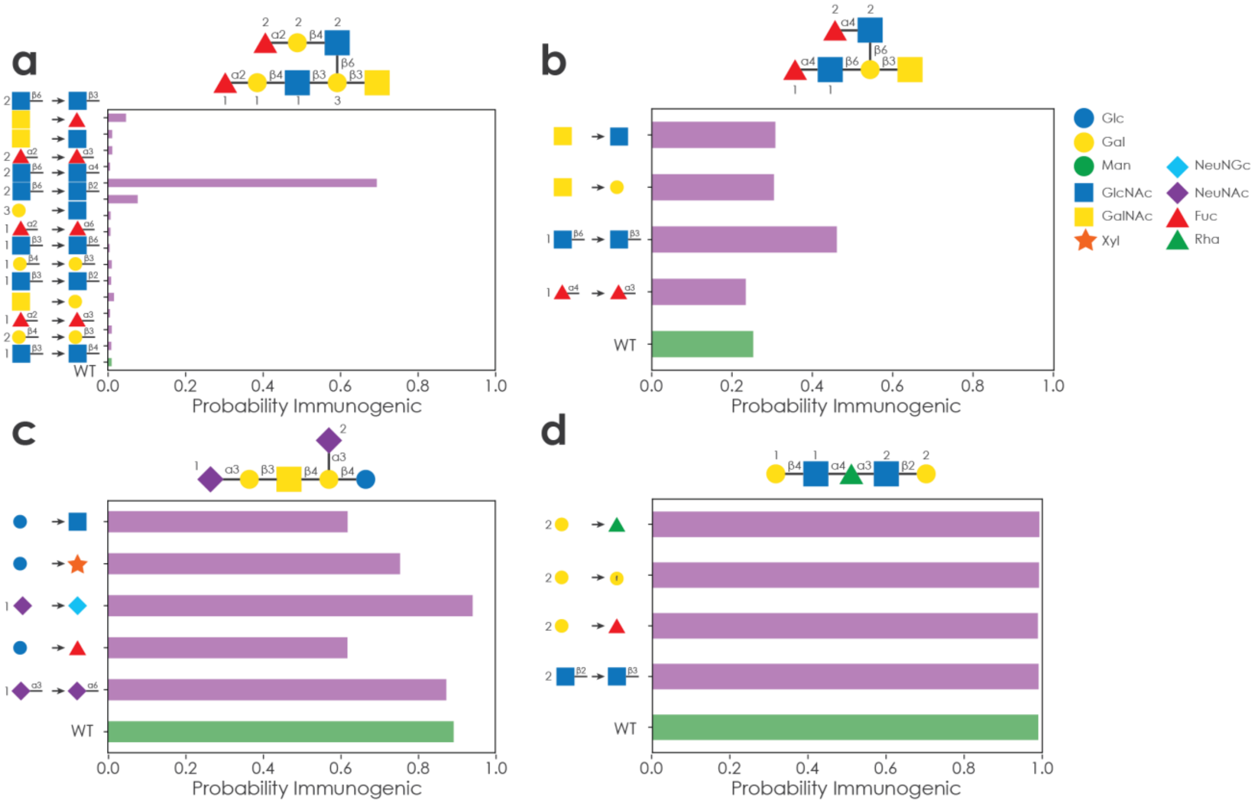


**Supplementary Fig. 10: Glycoword *in silico* modification to probe immunogenicity classifier.** (a-d) For 4000 iterations, a random position in a glycan was exchanged with a random monosaccharide or bond (depending on the position). The altered glycan was only retained and processed if the resulting glycowords existed in our dataset. Then, analogous to the masking experiments in Supplementary Fig. 9, we used our trained immunogenicity classifier for inference and plotted the predicted probability of the modified glycans to be immunogenic, with the wildtype glycan always located at the bottom. In case of ambiguity, a number indicates which monosaccharide was modified. Glycans are drawn in accordance with the symbol nomenclature for glycans (SNFG). The addition of an ‘f’ indicates the furanose form of the respective monosaccharide.

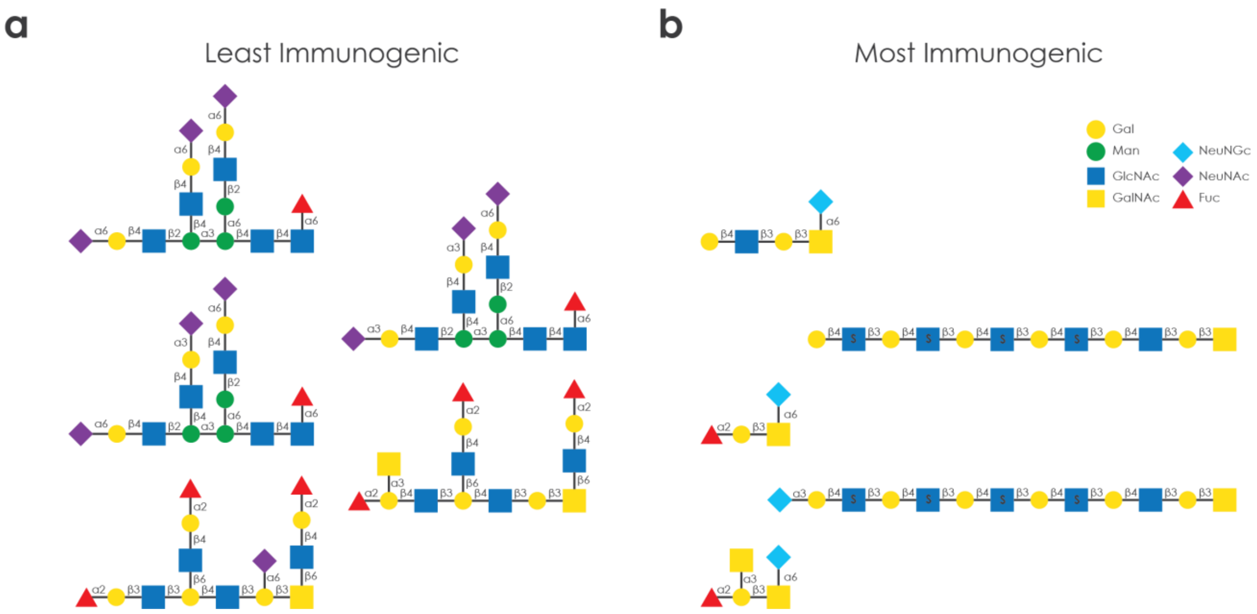


**Supplementary Fig. 11: Immunogenicity predictions of *Sus scrofa* glycans.** All glycans annotated with the species *Sus scrofa* were used as inputs for the trained immunogenicity classifier. Then, glycans were sorted by inferred human immunogenicity and the (**a**) five least immunogenic and (**b**) five most immunogenic glycans are shown here, respectively. Non-immunogenic glycans show high similarity to human glycans, with N-acetylneuraminic acid- (NeuNAc) and fucose-capped glycans. Porcine glycans predicted to be immunogenic, rich in N-glycolylneuraminic acid (NeuNGc) as well as sulfurylated monosaccharides, are supported by previous reports[22,23]. The full list of immunogenicity predictions can be found in Supplementary Table 12. Glycans are drawn in accordance with the symbol nomenclature for glycans (SNFG). The addition of an 'S' implies a sulfurylated monosaccharide.

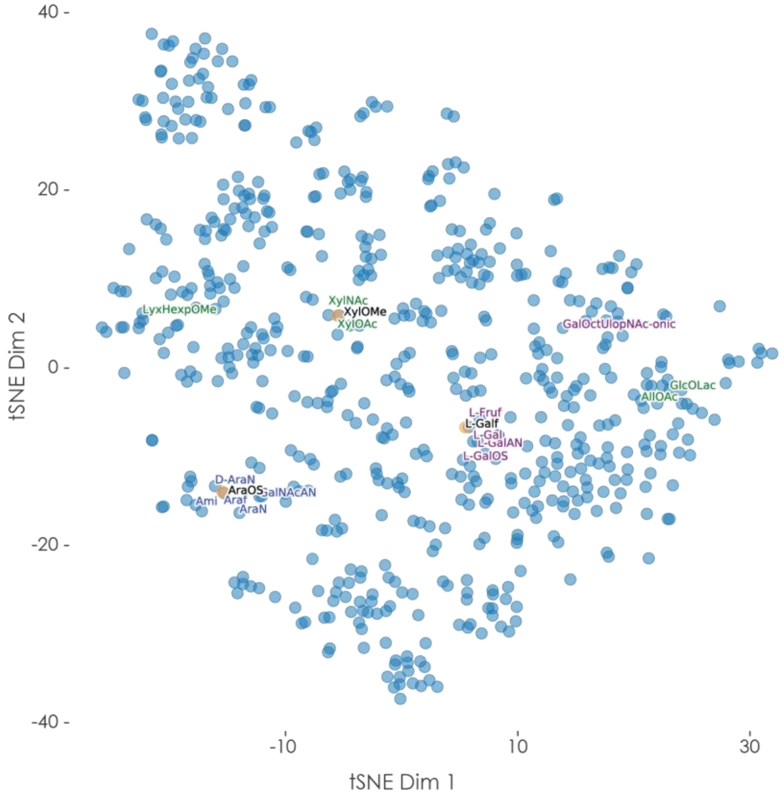


**Supplementary Fig. 12: Constructing embeddings for new glycoletters.** Embeddings for three potential glycoletters not observed in our dataset (O-sulfurylated arabinose (AraOS), O-methylated xylose (XylOMe), and L-galactofuranose (L-Galf)) were constructed by averaging their constituent character embeddings from the language model pre-training. Existing and artificial glycoletter embeddings were dimensionality-reduced via t-distributed stochastic neighbor embedding (t-SNE). Artificial glycoletter embeddings are colored orange, while observed glycoletters are colored blue. We then searched for existing glycoletter embeddings with the highest cosine similarity to the artificial glycoletter embeddings. For all artificial glycoletter embeddings, the five observed glycoletter embeddings with the highest cosine similarity are annotated and colored (AraOS: blue, XylOMe: green, and L-Galf: purple).
